# Melon ethylene-mediated transcriptome and methylome dynamics provide insights to volatile production

**DOI:** 10.1101/2020.01.28.923284

**Authors:** Ari Feder, Chen Jiao, Navot Galpaz, Julia Vrebalov, Yimin Xu, Vitaly Portnoy, Galil Tzuri, Itay Gonda, Joseph Burger, Amit Gur, Yaakov Tadmor, Arthur A. Schaffer, Efraim Lewinsohn, Nurit Katzir, Zhangjun Fei, James J. Giovannoni

**Author notes:** These authors contributed equally (Ari Feder, Chen Jiao).

## Abstract

During climacteric ripening large-scale transcriptional modifications are governed by ethylene. While ripening-related chromatin modifications are also known to occur, a direct connection between these factors has not been demonstrated. We characterized ethylene-mediated transcriptome modification, genome methylation dynamics, and their relation to organoleptic modifications during fruit ripening in the climacteric melon and an ethylene repressed line where the fruit-specific *ACC oxidase 1* (*ACO1*) gene was targeted by antisense. The *ACO1* antisense line exhibited mainly reduced transcriptional repression of ripening-related genes associated with DNA CHH hypomethylation at the onset of ripening. Additionally, transcription of a small set of ethylene-induced genes, including known ripening-associated genes, was inhibited by ACO1 repression and this inhibition was associated with CG hypermethylation. In the *ACO1* antisense line, the accumulation of aromatic compounds, which are mainly derived from the catabolism of amino acids, is known to be inhibited. One of the ethylene-mediated transcriptionally up-regulated genes, *CmTHA1*, encoding a threonine aldolase, exhibited differential cytosine methylation. Threonine aldolase catalyzes the conversion of L-threonine/L-allo threonine to glycine and acetaldehyde and thus is likely involved in threonine-dependent ethyl ester biosynthesis. Yeast mutant complementation and incubation of melon discs with labeled threonine verified CmTHA1 threonine aldolase activity, revealing an additional ethylene-dependent amino acid catabolism branch involved in climacteric melon ripening.

## Introduction

Ethylene, widely known as the ripening hormone, underlines climacteric fruit ripening. Widely studied climacteric fruits include tomato and melon where following seed maturation ripening competence is achieved and a burst of autocatalytic ethylene orchestrates coordinated changes in fruit pigmentation, aroma, cell wall modifications and tissue softening (Giovannoni, 2004; Yano and Ezura, 2016). This transition is accompanied by reprograming of expression of thousands of genes (Alba et al., 2005; Saladié et al., 2015; Shin et al., 2017; Yano et al., 2018). Fruit ethylene biosynthesis is dependent on the activity of multiple genes including transcription factors (TFs) and changes in histone-mediated repression of MADS-box and/or NAC domain TFs (Giovannoni et al., 2017; Rios et al., 2017; Lu et al., 2018). From a biochemical flux perspective autocatalytic ethylene involves transcriptional up-regulation of *cystathionine-γ-synthase* (*CGS*), the first committed step in synthesis of methionine, the amino acid precursor of ethylene (Alba et al., 2005). Much of our knowledge of climacteric fleshy fruit ripening derives from studies in tomato. Melon is also widely studied as it exhibits extreme genotypic and phenotypic variation including in climacterism (Burger et al., 2010; Gur et al., 2017; Galpaz et al., 2018). In contrast to tomato, fruit carotenoid accumulation is ethylene independent in melon and controlled by a ‘golden’ SNP in the *CmOr* gene (Tzuri et al., 2015). Introduction of a ‘golden’ SNP harboring allele into tomato significantly elevates fruit nutritional value through increased carotenoid accumulation (Yuan et al., 2015; Yazdani et al., 2019).

The *ACO1* antisense ethylene deficient mutant has proven an important tool in the study of climacteric melon fruit ripening (Ayub et al., 1996; Pech et al., 2008). In particular, *ACO1* antisense fruits fail to degrade chlorophyll and retain green rind color. The stay-green (SGR) phenotype is mediated by reduced activity of the ethylene-regulated SGR protein resulting in reduced chlorophyll degradation (Alba et al., 2005; Barry et al., 2008; Shimoda et al., 2016). *ACO1* antisense also results in greatly reduced fruit aromatic volatile compounds including acetate esters and ethyl esters (Homatidou et al., 1992; Bauchot et al., 1998; Flores et al., 2002). During melon ripening, aroma is largely attributed to ester biosynthesis involving the conversion of aldehydes into alcohols by alcohol dehydrogenases (ADHs) (Manríquez et al., 2006; Jin et al., 2016) followed by activity of alcohol acyltransferases (AATs) (Shalit et al., 2001; El-Yahyaoui et al., 2002; El-Sharkawy et al., 2005). Several amino acid metabolism pathways and associated ethylene-regulated genes underlying ester biosynthesis have been identified in melon. These include metabolism of branched-chain amino acids, methionine and phenylalanine (Wyllie et al., 1995; El-Yahyaoui et al., 2002; El-Sharkawy et al., 2005; Gonda et al., 2010; Gonda et al., 2018). Acetaldehyde is synthesized from pyruvate, the end product of glycolysis via pyruvate decarboxylase (PDC) (Tietel et al., 2011). Recently CmPDC1, a ripening induced melon pyruvate decarboxylase has been shown to be involved in acetaldehyde biosynthesis and downstream ester accumulation (Wang et al., 2019). Acetaldehyde has been shown to be a limiting factor in ethanol and derived ester levels in feijoa and strawberry (Pesis and Avissar, 1990; Pesis et al., 1991). Similar observations were made in apple for different aldehydes (De Pooter et al., 1983). Goulao and Oliveira (2007) hypothesized that threonine aldolase activity might also serve as a limiting factor in acetaldehyde-derived volatile biosynthesis during apple ripening based on transcriptional up-regulation of a putative L-allo-threonine aldolase gene. Due to their contributions to fleshy fruit quality, there is growing interest in biosynthesis of aromatic compounds, though limited variance is observed in modern cultivars of some important species such as tomato (Tieman et al., 2017).

Recently fleshy fruit development has been shown to be dependent upon chromatin remodeling. For example, during early tomato fruit and floral development, *SlFIE* and *SlEZ1*, members of the polycomb repressive 2 (PRC2) complex, play critical roles in fruit development as demonstrated by their repression (How Kit et al., 2010; Liu et al., 2012; Bucher et al., 2018). The PRC2 complex, which belongs to the polycomb group (PcG), catalyzes trimethylation of histone H3 lysine 27 (H3K27me3), negatively regulating gene expression (Holec and Berger, 2012). Additional regulators found in plants include histone variants. Histone H3.1 is enriched in silent genomic regions enriched with both H3K27me3 and DNA methylation, negatively correlating with gene expression; in contrast, H3.3 associates with actively transcribed genes in *Arabidopsis* (Stroud et al., 2012). DNA methyltransferase inhibition promotes premature ripening in tomato, suggesting an active remodeling mechanism prevents premature ripening through DNA methylation (Zhong et al., 2013). During tomato fruit ripening, *SlDML2*, a DEMETER-like DNA demethylase is transcriptionally up-regulated, and its repression or gene editing resulted in genome-wide hypermethylation, including in the promoters of ripening transcription factors resulting in inhibition of ethylene biosynthesis and ripening repression (Liu et al., 2015; Lang et al., 2017). Further genetic evidence of an active gene suppression mechanism during early fruit developmental stages was demonstrated through manipulation of the PRC2 member *SlMSI1* (Liu et al., 2016).

DNA CHH methylation typical of heterochromatic transposable elements (TEs) is significantly increased in fruits. In addition CHH methylation exhibits dynamics during fruit development and is affected by ripening mutants and has been associated with gene expression changes (Zhong et al., 2013; Corem et al., 2018; Lu et al., 2018). An example of interaction among multiple chromatin remodeling components was demonstrated by a mutation in the rice CHH methyltranferase gene *OsDRM2*, resulting in the loss of H3K27me3 and de-repression of genes (Zhou et al., 2016). In contrast to chromatin regulation of the ripening transition and ethylene biosynthesis (Giovannoni et al., 2017; Lu et al., 2018), the downstream involvement of chromatin dynamics in the ethylene response of climacteric fruits remains poorly understood.

To test whether there is a direct link between ethylene-regulated genes and chromatin dynamics we performed transcriptome and methylome comparisons of wild-type Védrantais (VED) and *ACO1* repressed melon fruit during early ripening and discovered ethylene-dependent methylome dynamics associated with ripening gene expression. A subset of this data suggested a role of threonine aldolase (CmTHA1) in melon ripening and fruit quality. Functional analysis confirmed that CmTHA1 plays a role in plant secondary metabolism, specifically production of volatile compounds integral to melon fruit quality.

## Results

To determine the effect of ethylene on transcriptional regulation during melon fruit development, RNA-Seq transcriptome profiling was performed on mesocarp tissue at five stages: Half Size (HS), Full Size (FS), Onset of Ripening (OR), Early Climacteric (EC), and Full Climacteric (FC; Fig. 1A, Supplemental Tables S1-S3). Significant ethylene accumulation in VED was detected at the EC and FC stages, while the *ACO1* antisense line exhibited >99% reduction in ethylene evolution (Fig. 1B). The analysis confirmed abundant mRNA accumulation of *ACO1* in VED (*MELO3C014437*; http://cucurbitgenomics.org/), correlating with ethylene accumulation, while in the *ACO1* antisense line *ACO1* mRNA was dramatically reduced (Fig. 1C). Ethylene accumulation in VED at EC was accompanied by differential expression of 2,687 genes when compared to *ACO1* antisense (adjusted p<0.05, minimum 2-fold difference), of which 2,070 were up-regulated and 617 were down-regulated in *ACO1* compared to VED (Fig. 1D-E), indicating that the predominant ethylene effect during VED fruit ripening manifests as down-regulation of fruit gene expression.

**Figure 1:**
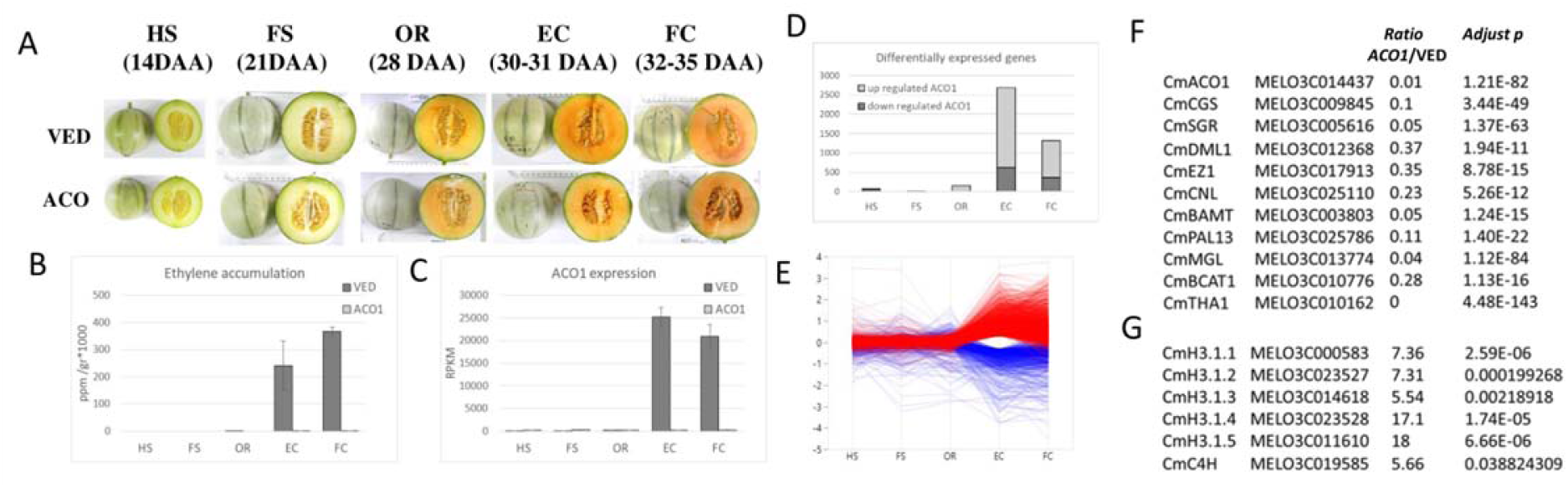
WT (VED) versus *ACO1* antisense melon development, ethylene synthesis and gene expression. **A.** Developmental stages in which fruits were harvested of WT cv. Vedrantais (VED) and the *ACC oxidase 1* antisence line (*ACO1).* HS: half size, FS: full size, OR: onset of ripening, EC: early climacteric, FC: full climacteric, DAA: days after anthesis **B.** Ethylene accumulation during development (nl + gr^−1^ + h^−1^. **C**. RNAi target gene *(ACO1)* expression pattern during development (RPKM). **D.** Summary of differently expressed genes (DEGs) between VED and ACO1 (adj p<0,05, minimum 2 fold difference) during the five fruit developmental stages. **E.** Parallel plot of log RPKM (ACO/VED) of 2687 DEGs, from EC stage, during the five fruit developmental stages. In red ACO1 up regulated 2070 genes, in blue ACO1 down regulated 617. **F-G.** selected DEGs between VED and *ACO1.* Ratios represent the RPKM ratio of *ACO1/M*ED average expression at EC stage. **F.** genes which are up regulated by ethylene and inhibited in *ACO1*. **G.** Genes which are down regulated by ethylene while highly expressed in *ACO1.*

### Dynamics of chromatin remodeling factors

The role of DNA methylation in ethylene-mediated gene expression differences was of interest due to recent reports regarding the role of DNA methylation and chromatin remodeling during fruit ripening (Zhong et al., 2013; Liu et al., 2015; Lang et al., 2017; Lu et al., 2018). DNA bisulfite sequencing was performed to determine cytosine (C) methylation in EC fruit from both genotypes (Supplemental Tables S4-S6). Similar to previous reports in Arabidopsis (Zhang et al., 2006), hypermethylated sites in melon were also found mainly in transposable element (TE)-rich heterochromatic regions (Fig. 2A). Total cytosine methylation in the EC tissue of VED was 23%, while the *ACO1* mutant exhibited 4% less genome-wide methylation (19%) in all cytosine context (Supplemental Table S6). This hypomethylation included regions associated with coding sequences and again in all cytosine contexts, and generally within a window spanning 2 kb up- and downstream of coding sequence (Fig. 2B-D). Methylome analysis between VED and ACO1 antisense EC fruit revealed 52,426 differentially methylated regions (DMRs) in the CHH context and 9,866 in the CG context (Fig. 2A, Supplemental Table S7). 46.5% of the CHH and 38% of the CG DMRs were found to associate with gene regions defined as +/-2 kb from the ends of coding sequences. Total methylation of gene-associated DMRs was decreased by 27% in the *ACO1* antisense line compared to VED (calculated by the sum of average methylation in each site multiplied by total DMR length), Suggesting ethylene regulates active CHH methylation leading to transcriptional repression. Of the 2,687 DEGs at the EC stage between VED and *ACO1* antisense, 1,988 (74%) were associated with DMRs (Supplemental Table S7). We noted a general tendency toward hypomethylation of genes not repressed by ethylene in *ACO1* antisense fruit (Fig. 2E-F), most prevalently in the CHH context.

**Figure 2:**
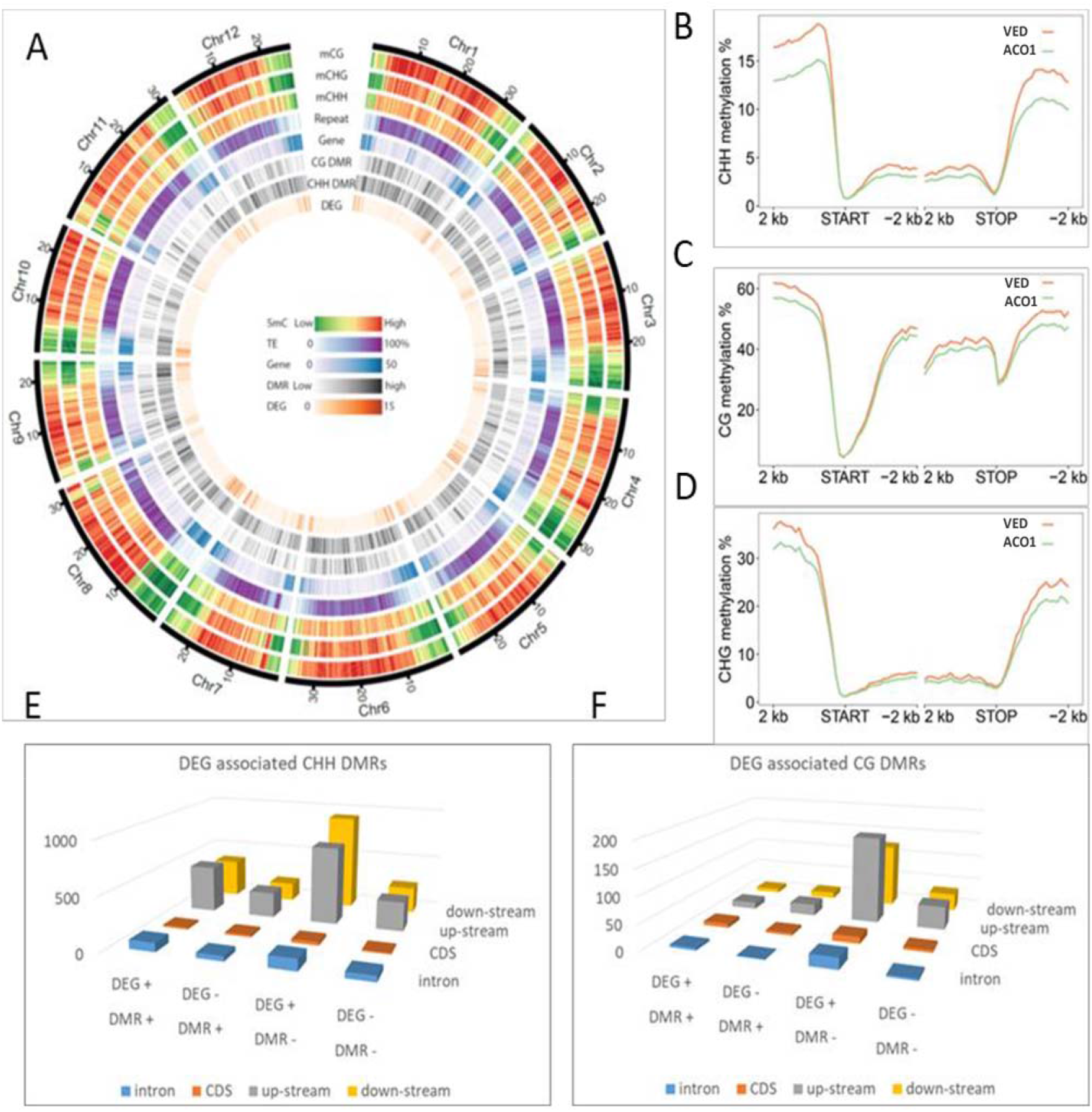
Genomic DNA methylation and it’s association to gene coding regions and differentially expressed genes (DEGs) **A.** Density plot of: 5-methylcytosine in sequence contexts (mCG, mCHG, mCHH), transposable elements (repeat), gene coding regions, CG DMRs, CHH DMRs, and DEGs. **B-D.** Coding gene associated DNA methylation in 2Kb windows upstream to start and downstream to stop codons, in the CHH, CG, CHG context respectively, on average of all coding sequences, in VED (orange), and the AC01 antisence line (green). **E-F.** Association between DMRs and DEGs. DEG (-): gene transcripts which are down regulated in ACO1, DEG (+): up regulated in ACO1. DMR (-): Hypomethylted regions in the ACO1, DMR (+) Hyper methylated regions in ACO1, in the different transcript context of: upstream to start codon, down stream to stop codon, introns, or coding sequence (CDS).

Although most of the genes in this this study consist of up-regulated DEGs (as compared to VED) associated with hypomethyated DMRs in the EC tissue of *ACO1* antisense in the CG or CHH context, 30 DEGs did show CG hypermethylation associated with transcriptional down-regulation in the *ACO1* antisence. These genes include: *CmSGR* (*MELO3C005616*), *CmCGS* (*MELO3C009845*), and *CmEZ1* (*MELO3C017913*), the ortholog of tomato *SlEZ1* involved in H3K27me3-mediated gene silencing (Table S8; How kit et al 2010) and could be targets of DNA demethylase activity. *SlDML2*-dependent hypomethylation is the main mechanism underlying DNA methylation dynamics during fruit ripening characterized to date in tomato (Liu et al., 2015; Lang et al., 2017). Melon climacteric ripening is also associated with increased expression of a DNA glycosylase/demethylase, *CmDML1* whose expression is reduced to 37% of WT with *ACO1* repression (*MELO3C012368.2*, Fig. 1F, Supplemental Fig. S1). Unlike the tomato ripening-upregulated *SlDML2* gene which is an ortholog of Arabidopsis *REPRESSOR OF SILENCING1 (ROS1)*, *CmDML1* is the ortholog of Arabidopsis *DEMETER* (*DME*) (Zemach et al., 2010; Zhong et al., 2013). It is especially noteworthy that loss of *AtDME* function can occur with some loci still becoming hypomethylated, likely due to RNAi-mediated DNA methylation dynamics (Hsieh et al., 2009; Zemach et al., 2010). As such, while decreased *ACO1* genome methylation is not consistent with the observed reduction of *CmDML1* expression in the antisense line, alternative methylation/demethylation mechanisms likely contribute to maintaining relative hypomethylation in the absence of ethylene. Likely candidate genes contributing to this phenomena (e.g methyltransferases) were not revealed by our transcriptome analysis but the reduction in CmCGS necessary for downstream S-adenosyl-methionine (SAM) production (the methyl donor for DNA methylation) may provide a general mechanism for reduced methylation in *ACO1* antisense fruit.

Ethylene also influences histone variant transcriptional dynamics which could further influence gene expression. All five H3.1 histone variants found in melon (*MELO3C000583*, *MELO3C023527*, *MELO3C014618*, *MELO3C023528*, and *MELO3C011610*, Fig. S2) displayed ethylene-dependent down-regulation in VED, which is arrested in *ACO1* antisense fruit, resulting in a 5-18 fold expression increase in their relative mRNA abundances in the *ACO1* antisense line as compared to VED (Fig. 1G). The increased abundance of H3.1 histones which are generally associated with gene silencing could also be responsible for reduced expression of some genes repressed in *ACO1* ethylene reduced fruit.

### Genetic regulation of aromatic compound accumulation

Volatiles are a significant component of consumer appreciation of ripe fruit (Gonda et al., 2016). Amino acid catabolism results in the production of a wide array of volatiles in melon. L-phenylalanine derived volatiles represent an important group of melon volatiles, and key pathway genes are both regulated by ethylene and have associated DMRs revealed in this study. For example, *CmCNL* (*MELO3C025110*) encoding an (E)-cinnamic acid:coenzyme A ligase involved in (E)-cinnamaldehyde biosynthesis, and *CmBAMT* (*MELO3C003803*) encoding a benzoic acid:S-adenosyl-L-methionine carboxyl methyltransferase involved in methyl benzoate biosynthesis (Gonda et al., 2018), exhibited strong ethylene-dependent up-regulation (Fig. 1F) in addition to DMRs in both the CG and CHH contexts associated with *CmBAMT* (Supplemental Table S7). Two additional genes, *phenylalanine ammomia-lyase 13* (*CmPAL13*, *MELO3C025786*) and *cinnamate 4-hydroxylase 1* (*CmC4H1*, *MELO3C019585*), previously suggested to be involved in melon rind phenypropanoid synthesis (Feder et al., 2015), were also regulated by ethylene. *CmPAL13* was up-regulated by ethylene, while *CmC4H1* mRNA was down-regulated (Figs. 1F,G), suggesting the involvement of these genes in ethylene-mediated metabolic alternations resulting in channeling flux toward cinnamic acid and downstream volatiles in the maturing fruit. *CmC4H1* associated with a CHH DMR, while *CmPAL13* associated with DMRs in both the CHH and CG contexts (Supplemental Table S7). L-methionine-derived volatiles are dependent upon the *L-meththionine-γ-lyase* (*CmMGL*, *MELO3C013774*), which is involved in ethyl ester biosynthesis through L-isoleucine (Gonda et al., 2013). In addition, branched-chain amino catabolism in melon derive from α-keto acids of L-isoleucine, L-leucine and L-valine, through the activity of the enzyme encoded by *CmBCAT1* (*MELO3C010776*), a branched-chain amino acid transaminase which is highly expressed in climacteric ripening melon fruits (Gonda et al., 2010). These two genes, *CmMGL* and *CmBCAT1*, also exhibited strong ethylene-dependent transcriptional up-regulation during VED fruit ripening (Fig. 1F) and a CHH DMR is associated with *CmMGL* expression changes (Supplemental Table S7).

### Functional validation of *CmTHA1*’s role in melon fruit volatile synthesis

L-threonine contributes to volatile biosynthesis via L-isoleucine, which is catabolized into esters (Gonda et al., 2013). Here we show that ethylene regulates *CmTHA1* (*MELO3C010162*, Fig. 1F), which encodes a putative L-allo threonine aldolase. *CmTHA1* associated with four DMRs in both the CHH and CG contexts (Supplemental Table S7). The melon genome harbors two homologs to this gene, *CmTHA2* (*MELO3C017520*) and *CmTHA3* (*MELO3C004421*). *CmTHA1* displayed the greatest mRNA accumulation in EC fruit of VED and was the only one significantly affected by ethylene, suggesting its involvement in ripening (Supplemental Fig. S3). Functional assessment of *CmTHA1* was performed via yeast complementation in a similar manner as previously described (Jander et al., 2004), using *∆gly1*, a yeast deletion mutant in strain BY4741 (Giaever et al., 2002). Gly1 is a low specificity threonine aldolase catalyzing the cleavage of both L-threonine and L-allo-threonine to glycine (Liu et al., 1997). *∆gly1* was transformed independently with vector pESC-URA (*∆gly1*-URA) and with the same vector harboring *CmTHA1* coding sequence under the galactose inducible Gal1 promoter (*Δgly*1-*URA*-*CmTHA*). Transformed strains were verified by PCR (Fig. 3A). Confirmed strains were grown on yeast extract– peptone–dextrose (YPD) control medium (Fig. 3B) along with different dropout media. Growth upon uracil dropout medium verified the ability of the pESC-URA vector to facilitate recovery from uracil auxotrophy (Fig. 3C). Glycine dropout medium supplemented with glucose verified that *∆gly1* is indeed a glycine auxotroph as well as the two *∆gly1*-URA and *Δgly*1-*URA*-*CmTHA* transformants (Fig. 3D). Changing glucose to galactose, allowing Gal1 promoter activation, confirmed that CmTHA1 could relieve *∆gly1* glycine auxotrophy (Fig. 3E), indicating that CmTHA1 harbors THA activity.

**Figure 3:**
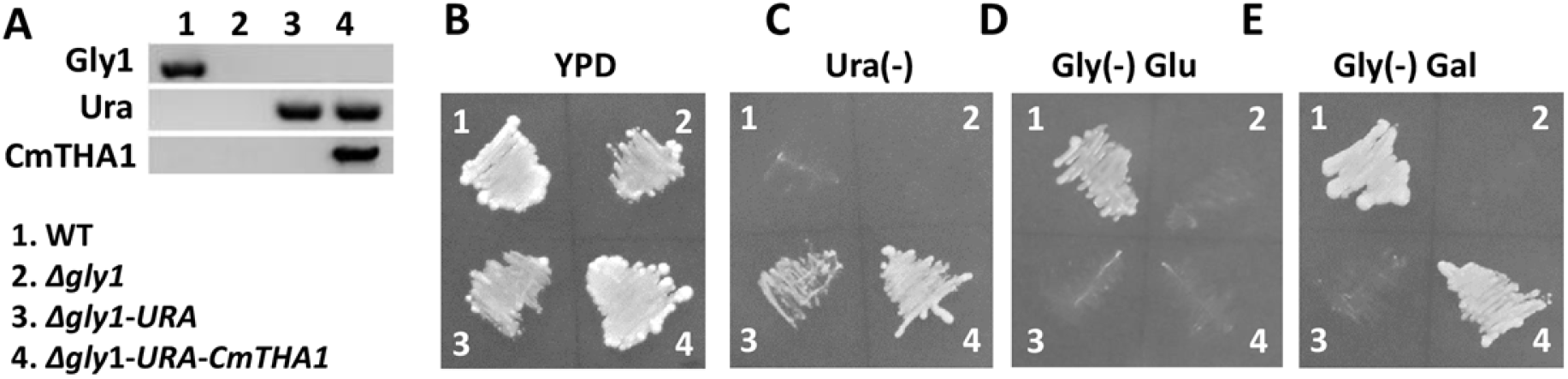
Functional analysis of *CmTHA1* in yeast. **A.** Agarose gel of yeast DNA extraction PCR, performed upon: 1. WT yeast strain 2. Δ*gly1* threonine aldolase deletion mutant. 3. Δ*gly1* transformed with pESC-URA plasmid, 4. Δ*gly1* transformed with the pESC-URA plasmid, added with *CmTHA1* under GAL1 promoter. Each strain checked with three sets of primers to the genes: Gly1-yeast endogenous threonine aldolase, Ura-plasmid 5’-phosphate decarboxylase *(URA3), CmTHA1-* melon threonine aldolase 1. **B.** 1-4 yeast strains as in A grown on YPD medium. **C.** strains grown on uracil dropout medium. **D.** strains grown on glycine dropout medium supplemented with glucose. **E.** strains grown on glycine dropout medium supplemented with galactose.

To test the threonine aldolase catalytic activity in melon, fruit disks were incubated with ^13^C_4_,^15^N-L-threonine, followed by GC-MS analysis. Two types of labeling were detected. The +2m/z labeling was detected in acetaldehyde, confirming aldolase activity. Similar +2 m/z labeling was also detected in ethanol, and the ethyl esters (ethyl acetate, ethyl butanoate, ethyl (methylthio) acetate, ethyl 2-methylbutyrate, and ethyl propanoate) (Fig. 4). In addition, stronger +3 m/z labeling was observed for ethyl 2-methybutyrate and ethyl propanoate, both of which were previously observed to be labeled similarity after incubation with ^13^C_6_-L-isolecine and positioned in the aldehyde derived moiety (Fig. 4C-D) (Gonda et al., 2013). Relatively low labeling of the +2 m/z fraction was anticipated to occur due to higher affinity of CmTHA1 to L-allo threonine and/or availability of acetaldehyde biosynthesized from pyruvate. The ester +2 m/z labeling position by the mass fragments of ethyl propanoate is a clear indication of its derivation from ethanol (Fig. 4E-F).

**Figure 4:**
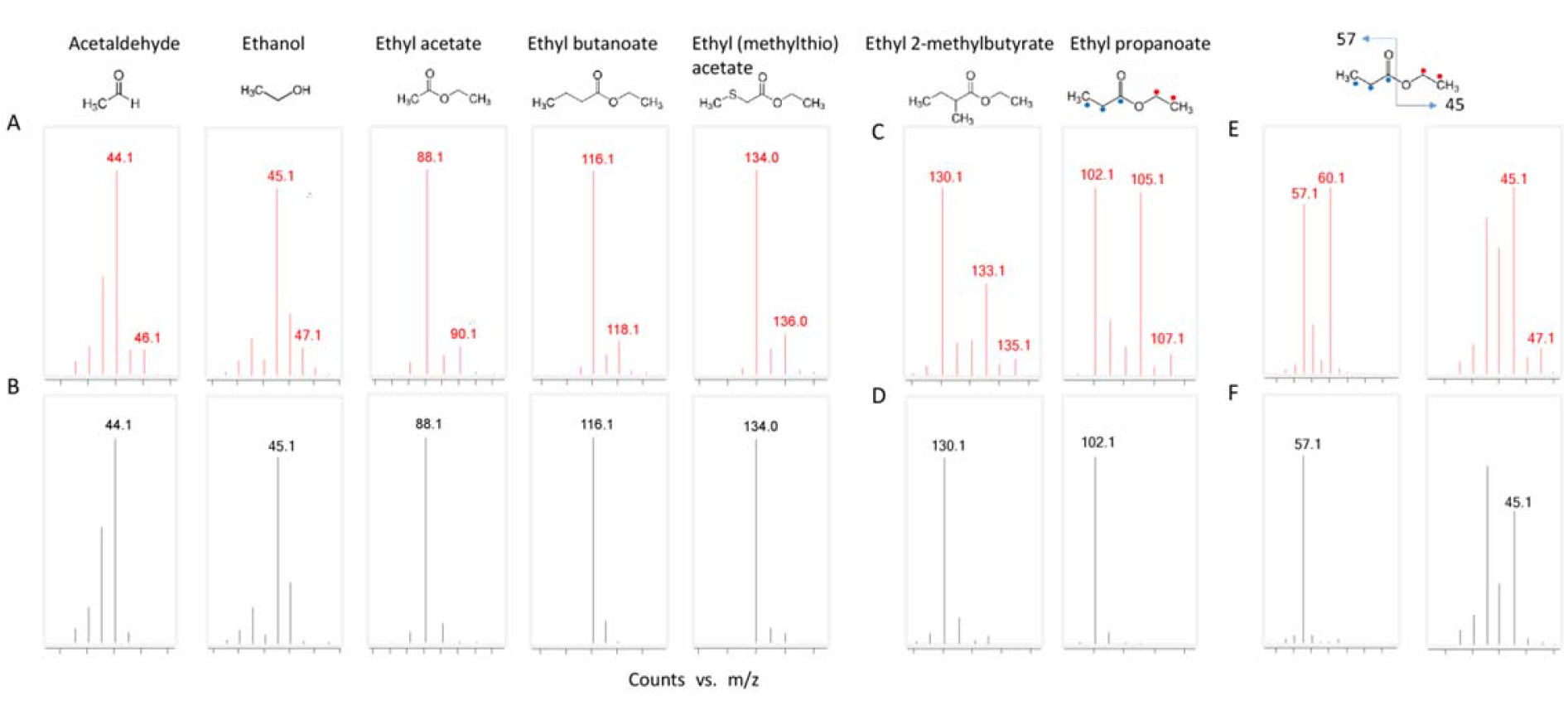
Mass spectra of volatiles isolated from melon fruit disks incubated with ^13^C4-L-Threonine. A, C, E (in red) are labeled treatments with corresponding unlabeled controls below in black (B, D, F). A & B: compounds enriched with +2 m/z labeling. C & D: compounds enriched with both +3 m/z and +2m/z labeling. +3 m/z labeling of these compounds was previously reported after incubation with L-isoleucine. Position of+3 m/z labeling is marked by blue dots (Gonda et al 2013). E & F mass fragmentation of ethyl propanoate.

## Discussion

### Climacteric melon ripening transcriptome activity shifts toward down-regulation of gene expression via increased gene methylation

Changes in DNA methylation have been implicated in ripening control in tomato (Zhong et al., 2013), a climacteric fruit whose ripening is dependent upon ethylene. In general tomato ripening occurs in the context of demethylation of ripening gene promoters at regions at or adjacent to transcription factor binding sites. The role of ethylene in genome methylation changes has not been specifically examined in prior studies. Here we show the effect of ethylene upon DNA methylation through examination of climacteric (VED) and non-climacteric melon where the latter is achieved via antisense repression of the ethylene synthesis gene *ACO1* (Ayub et al., 1996). Chromatin remodeling factor genes are among those responding to ethylene in this system, suggesting a role for ethylene-mediated histone changes during melon ripening (Fig. 1G). Chromatin methylation during ripening in VED was also recently reported at rates of 75%, 63% and 91% hypomethylation in the CG, CHG and CHH contexts, respectively, in DMRs during the VED transition from unripe to ripe (Lu et al., 2018). The same report analyzed two non-climacteric varieties that trended toward hypermethylation of DMRs in the CHH context during the immature and unripe to fully ripe transition. It is important to note that Lu et al. (2018) specifically compared unripe (20 DAA) and ripe (40 DAA) VED fruit, spanning a developmental transition encompassing both ethylene-dependent and independent physiological and biochemical changes. The long duration between compared stages precluded any direct correlation with the climacteric or ripening ethylene induction in VED. Changes in sugars, carotenoids, organic acids and part of cell wall metabolism all occur in this period and are known to be ethylene independent in melon (Ayub et al., 1996; Pech et al., 2008), and their relationship to chromatin dynamics remain unclear. Neither has any direct relationship between the plant hormone ethylene and methylome changes been established even though ethylene is a likely candidate for mediating such ripening-related changes in climacteric fruit. Here we characterized methylome changes in identically aged fruit at the early ripening stage, EC (30-31 DAA) in WT and ethylene repressed fruit to address whether ethylene is involved in the methylation dynamics of ripening genes. We observed that ethylene-mediated methylome changes are substantial and the main effect on DNA methylation associated with ethylene in WT climacteric melon fruit ripening is hypermethylation.

Ethylene mainly caused down-regulation of genes which are also associated with CHH hypermethylation in WT (Fig. 1D). In addition, 30 WT ethylene induced genes are associated with CG hypomethylation, suggesting possible DNA demethylase involvement. This possibility was supported by the observation of ethylene-dependent up-regulation of *CmDML1* (Fig.1F), the orthologue of Arabidopsis *DME*. Tomato ripening occurs with transcriptional up-regulation of a DNA demethylase, *SlDML2*, the orthologue of Arabidopsis *ROS1*. This observation is not necessarily surprising in the context of recently reported evidence of convergent evolution of different regulatory targets in the evolution of fruit ripening (Lu et al., 2018). AtDME functions in the CG hypomethylation-dependent activation of gene expression (Hsieh et al., 2009). We show ethylene-dependent transcriptional up-regulation of *CmDME1* associated with a corresponding decrease in CG methylation and transcriptional up-regulation of *CmSGR*, *CmCGS*, and *CmEZ1*, suggesting that these genes are possible CmDME1 targets.

Further investigation into possible DNA methylation regulators based on melon fruit gene expression changes suggests possible CmCGS involvement. In tomato, ethylene-regulated transcriptional up-regulation of *cystathionine-γ-synthase* (*CGS*), the first committed step in methionine biosynthesis, controls the biochemical flux toward ethylene and is part of the ethylene autocatalytic mechanism (Alba et al., 2005). From methionine, ethylene is synthesized through S-adenosyl-L-methionine (SAM) which also serves as the main methyl donor for cytosine methylation catalyzed by DNA methyltransferases. Disruption of this pathway, including at SAM biosynthesis in Arabidopsis, results in genome hypomethylation leading to de-repression of TEs and activation of gene expression (Rocha et al., 2005; Groth et al., 2016; Meng et al., 2018; Yan et al., 2019). We demonstrate CmCGS also shows ethylene-dependent transcriptional up-regulation (Fig. 1F), and as such that *CGS*-dependent autocatalytic ethylene is conserved between tomato and melon. As an ethylene synthesis biochemical flux regulator, CmCGS is a limiting factor in SAM biosynthesis, consistent with the decreased capability of *ACO1* antisense to down-regulate gene expression (Fig. 1D-E) through cytosine methylation (Fig. 2) due to limited availability of the necessary methyl donor.

In Arabidopsis, H3.1 histone variants were shown to associate with silent areas of the genome (Stroud et al., 2012). The ethylene-dependent transcriptional down-regulation of all melon H3.1 variants (Fig. 1G, Supplemental Fig. S3) suggests the ethylene-dependent transcription activation (Fig. 1D) is additionally mediated at least in part by down-regulation of H3.1 histone variants.

### CmTHA1 contributes to melon fruit volatile production

Ethylene is known to regulate different amino acid catabolism pathways involved in fruit aroma including: L-methionine (via CmMGL), branched-chain amino acids (via CmBCAT1) and L-phenylalanine (via CmBAMT, CmCNL). In addition, responsiveness of *CmPAL13* and *CmC4H1* to ethylene suggested additional upstream flux channeling that may influence levels of these amino acids and their volatile metabolic products (Fig.1 F-G).

While the final steps in volatile ester biosynthesis involving AHD and AAT are largely understood, earlier steps in aldehyde formation remain uncertain. Recently CmPDC1 was found to mediate acetaldehyde biosynthesis in ripening melon fruit though additional factors are certainly involved (Wang et al., 2019). We demonstrate here that one such factor is CmTHA1 as demonstrated by +2 m/z labeling of acetaldehyde originating from threonine (Fig. 4). The resulting +2 m/z labeled compounds follow the general paradigm of ester biosynthesis in which acetaldehyde is converted to ethanol by ADH and subsequent AAT-dependent incorporation into ethyl esters (ethyl acetate, ethyl butanoate, ethyl (methylthio) acetate, ethyl 2-methylbutyrate, and ethyl propanoate).

### Summary

We investigated basic chromatin remodeling at the level of DNA cytosine methylation specifically as influenced by ethylene during fruit ripening through comparison of early ripening WT climacteric (VED) and ethylene repressed transgenic (*ACO1* antisense) melon fruit. It is well known that modern cantaloupe melon varieties produce reduced aromatic compounds as a consequence of breeding efforts focused on increased shelf life through limiting ethylene biosynthesis and/or perception (Aubert and Bourger, 2004; Obando-Ulloa et al., 2008). We demonstrate that melon fruit severely limited in ripening ethylene production have substantially altered gene expression at the initial ripening stage and that many genes altered in expression are associated with DMRs, the majority of which are associated with hypomethylation in ethylene-repressed fruit and elevated gene expression as compared to WT. A smaller subset of genes show elevated expression with lower methylated DMRs in WT, a phenomena also reported in tomato (Lang et al., 2017). As ethylene is known to be important in melon volatile synthesis we focused efforts on characterization of ethylene and DMR-associated genes contributing to volatile synthesis. We identified and demonstrated the function of CmTHA1, a previously uncharacterized L-allo threonine aldolase contributing to volatile ester synthesis. The more comprehensive picture of ethylene effects on gene expression resulting from this study should prove helpful in designing breeding strategies focused on ethylene-regulated ripening components such as volatile synthesis to target and reverse the negative effect of ethylene reduction on aroma.

## Materials and methods

### Plant material and ethylene measurements

Seeds of the *ACO1* antisense line (Ayub et al., 1996), along with Védrantais (VED) were kindly provided by Maria-Carmen Gomez-Jimenez, Plant Physiology department, University of Extremadura, Spain. Plants were grown in a randomized design in pots inside a greenhouse under standard conditions during spring 2013 at Newe Ya’ar Research Center. Flowers were tagged at anthesis. Fruits were collected during development according to Table S1. Ethylene emission was measured as previously described (Galpaz et al., 2018).

### RNA-Seq library preparation, sequencing and data analysis

RNA extraction, library preparation and sequencing were performed according to the methods described previously (Zhong et al., 2011; Feder et al., 2015). Two to four biological replicates were performed for each sample. Raw RNA-Seq reads were processed using Trimmomatic (Bolger et al., 2014) v0.36 to remove adaptor and low-quality sequences. The cleaned reads were then aligned to the ribosomal RNA database (https://www.arb-silva.de/) with bowtie (Langmead, 2010) v1.0.0 (parameter ‘-v 3’) to filter out rRNA reads. The final cleaned reads were aligned to the melon genome v3.5.1 (Diaz et al., 2015) using HISAT (Kim et al., 2015) allowing up to two mismatches. Raw counts for each gene were then derived from the alignments and normalized to reads per kilobase of transcript, per million mapped reads (RPKM). Differentially expressed genes between WT climacteric (VED) and ethylene repressed transgenic (ACO1 antisense) melon fruit at each of the five developmental stages, half size (HS), full size (FS), onset of ripening (OR), early climacteric (EC), and full climacteric (FC), were identified with the DESeq2 package (Love et al., 2014). Genes with adjusted *P* value <0.05 and fold-change ≥2 were defined as differentially expressed genes.

### Whole-genome bisulfite sequencing and data analysis

DNA for bisulfite sequencing was extracted from isolated nuclei following Zhong et al. (2013). Bisulfite library construction and sequencing were performed at the Roy J. Carver Biotechnology Center, University of Illinois, Urbana-Champaign. Raw bisulfite sequencing reads were first processed to collapse duplicated read pairs into unique read pairs. The resulting reads were then processed to remove adaptor and low-quality sequences using Trimmomatic v0.36. Two biological replicates were combined for the downstream analysis to obtain a better coverage of the genome. The method for the whole genome methylation analysis was same as described previously (Zhong et al., 2013). Briefly, before alignment, each base cytosine in the reads and the double-strand genome (G in reverse strand) was replaced with thymine (or A if reverse strand in the genome). The converted reads were aligned respectively to the two converted strands of the genome using bowtie allowing up to two mismatches, and reads aligned to multiple locations were excluded from the analysis. Alignments from the two strands were combined and the original read sequences in the alignments were recovered. Finally, methylation status of each cytosine in the melon genome was calculated on the basis of the alignments. A sliding-window approach with a 100-bp window sliding at 50-bp intervals was used to identify context-specific DMRs. Windows with fewer than 5, 4 and 20 sequenced cytosine sites (≥4× coverage) in the CG, CHG and CHH contexts, respectively, were discarded. For each window, a Kruskal–Wallis test was performed, and the *P* values were corrected using Benjamini-Hochberg method (Benjamini and Hochberg, 1995). Windows with corrected *P* value <0.05 were identified as DMRs and overlapping DMRs were concatenated.

### Yeast transformation

*∆gly1* yeast obtained from the knockout library in the BY4741 background (Giaever et al., 2002), along with pESC-URA, were kindly provided by Maya Schuldiner’s lab at Weizmann Institute of Science. Coding sequence of *CmTHA1* was obtained from cDNA of ripe fruit of Charentais melon, following PCR with the THA1-F/R primers. pESC-URA was linearized using SmaI/HindIII restriction enzymes, following insertion of the PCR product with NEBuilder (New England BioLabs). Growing media, including custom made Kaiser synthetic complete Gly(-) dropout, were obtained by Formedium. Primer sequences used in this study are listed in Supplemental Table S9.

### Labeled L-threonine feeding and CG-MS analysis

Plants were grown in winter 2019 in greenhouse at Newe Ya’ar. Ripe fruits were collected, left at room temperature for 48h. Fruit mesocarp discs (approximately 1g each) were incubated with 100mM L-threonine-^13^C_4_,^15^N (Sigma) for 12 hours, after which tissue was frozen in liquid nitrogen and ground. Sample preparation for GC-MS was performed according to Gonda et al 2013. Data analysis was performed with the MassHunter software (Agilent).

## Accession numbers

## Acknowledgments

We thank Eric Richards for the insightful discussions. This research was supported The United States-Israel Binational Agricultural Research and Development Fund, Grant Award No. 3012114 (ICOB program) from the NSF-BSF Joint Funding Program, National Science Foundation Plant Genome Program grant No. 1339287 and the USDA-ARS.

## Supporting information

Additional Supporting Information may be found in the online version of this article.

**Supplemental Figure S1**. Phylogenetic tree of the DML glycosylase gene family.

**Supplemental Figure S2**. Phylogenetic tree of H3.1 and H3.3 histone variants.

**Supplemental Figure S3**. Melon threonine aldolase fruit gene expression.

**Supplemental Table S1**. Fruit sampling, measurements, RNA-Seq preparation and statistics.

**Supplemental Table S2**. RNA-Seq sample correlation.

**Supplemental Table S3**. RNA-Seq gene expression (RPKM).

**Supplemental Table S4**. DNA bisulfite sequencing statistics.

**Supplemental Table S5**. DNA bisulfite coverage percentage.

**Supplemental Table S6**. DNA methylation percentage.

**Supplemental Table S7**. Differentially methylated regions associated with protein-coding genes.

**Supplemental Table S8**. Putative CmDML1 targets.

**Supplemental Table S9**. Primers used in this study.

